# Early epigenetic priming of regeneration studied with a novel transgenic epigenetic reporter

**DOI:** 10.1101/2025.04.21.649771

**Authors:** Jian Ming Khor, Isabella Cisneros, Miranda Marvel, Aniket V. Gore, Kiyohito Taimatsu, Gennady Margolin, Daniel Castranova, Allison Goldstein, Ryan K. Dale, Brant M. Weinstein

## Abstract

Tissue regeneration requires previously differentiated cells to regain developmental plasticity. However, the upstream mechanisms initiating this process remain poorly understood. Here, we leverage a novel “EpiTag” transgenic zebrafish reporter line that enables real-time visualization of epigenetic silencing and activation to identify and carry out a comprehensive multi-omics analysis of cells undergoing epigenetic reprogramming during caudal fin regeneration. EpiTag GFP expression is transiently activated in cells contributing to regeneration between 12 to 16 hours post amputation (hpa), preceding the expression of canonical blastema markers. Single-cell RNA-seq reveals that GFP+ cells are restricted to regeneration-competent lineages such as pre-osteoblasts, proliferating cells, and wound epithelium. Integrated bulk RNA-seq, time-course RNA-seq, ATAC-seq, and bisulfite-seq on FACS-isolated GFP+ cells uncovers an early gene expression module enriched for chromatin regulators and a late gene expression module enriched for morphogenesis genes. Chromatin accessibility and DNA methylation changes are strongly associated with these late-expressed genes, suggesting epigenetic priming. We identify a number of epigenetic factors upregulated in the early gene expression module and show that ruvbl1 and ruvbl2, components of ATP-dependent chromatin remodeling complexes, are required for proper regeneration in both adult fins and larval tails. Our results establish EpiTag transgenics as a powerful *in vivo* tool for studying epigenetic reprogramming and highlight early chromatin remodeling events that enable activation of regenerative gene expression programs.

## Introduction

In regeneration-competent species such as zebrafish, adult tissues possess a remarkable capacity to rebuild complex structures following injury through reactivation of developmental gene expression programs (Beffagna, 2019; Gemberling et al., 2013). Zebrafish fin regeneration provides a superb example of this, acting as a useful and accessible model for studying how developmental reactivation is accomplished. Upon caudal fin amputation, differentiated cells at the wound site undergo dedifferentiation, migration, and proliferation to form the blastema, a highly proliferative structure that gives rise to new tissue (Pfefferli & Jaźwińska, 2015; Sehring & Weidinger, 2020). While significant advances have been made in elucidating the transcriptional and signaling networks that regulate dedifferentiation (Knopf et al., 2011; I. Sehring et al., 2022), epithelial-to-mesenchymal transition (EMT) (Klatt Shaw et al., 2021; Tang et al., 2022), tissue patterning (Quint et al., 2002), and cell proliferation (Grotek et al., 2013; Hui et al., 2022), the upstream mechanisms that initiate these regenerative processes remain unclear. Previous studies have hinted that changes in the epigenome may be a very early event required to trigger subsequent regenerative processes, including work showing that DNA methylation levels are reduced in the blastema during fin regeneration (Hirose et al., 2013), and that histone demethylase *kdm6b* is required for regenerative outgrowth (Stewart et al., 2009).

Epigenetic reprogramming, including changes in DNA methylation, histone modifications, and chromatin remodeling, plays a critical role in establishing and maintaining cellular identity during development (Cantone & Fisher, 2013). In contrast to development, tissue regeneration in adults requires previously differentiated cells to partially reverse their epigenetic state in order to re-acquire cellular plasticity and reactivate developmental programs (Katsuyama & Paro, 2011). Studies in zebrafish and other regenerative models have shown that regeneration is accompanied by the expression of genes normally silenced after embryogenesis (Brezitski et al., 2021; Karra et al., 2015; Weinberger et al., 2024). However, direct evidence for when and where early epigenetic changes occur during regeneration remains limited, in part due to a lack of tools to dynamically track epigenetic changes in real time in live animals, and to selectively profile epigenomic and transcriptomic changes in early regenerating cells.

To address this challenge, we leveraged the EpiTag reporter line, a novel transgenic zebrafish model that enables real-time, dynamic visualization of epigenetic silencing and activation in intact living animals at cellular-level resolution (Marvel et al., 2025). This reporter expresses a destabilized GFP (GFPd2) under the control of a CpG island from the *deleted in azoospermia-like (dazl)* gene linked to the *Xenopus ef1a* promoter. During normal development, the *dazl* CpG island and adjacent *Xenopus ef1a* promoter become rapidly methylated and EpiTag GFPd2 expression is silenced in all somatic tissues. However, during regeneration, EpiTag GFPd2 expression is turned on again, providing a sensitive readout of epigenetic activation. This system enables both high-resolution live imaging and easy isolation of cells undergoing epigenetic reprogramming during fin regeneration.

In this study, we combined the EpiTag reporter with a multi-omics approach, including single-cell RNA-seq, bulk RNA-seq, time course RNA-seq, ATAC-seq, and bisulfite sequencing, allowing us to precisely define the timing, cellular context, and molecular features of early epigenetic activation during zebrafish caudal fin regeneration. We show that the EpiTag reporter activity is transiently induced between 12 and 16 hours post amputation (hpa), preceding the expression of classical blastema markers such as *msx1b*. Single-cell analysis revealed that GFP+ cells are restricted to specific regenerative lineages, including pre-osteoblasts, proliferating epithelium, and wound epithelium. Integration of time-course RNA-seq with chromatin accessibility and DNA methylation profiling identified early-expressed gene modules enriched for RNA processing and chromatin regulators, followed by later modules associated with morphogenesis.

We identified a number of epigenetic factors up-regulated in blastema tissues early in the regenerative process, including *ruvbl1* and *ruvbl2*, conserved components of ATP-dependent chromatin remodeling complexes. We demonstrated through functional perturbation that both genes are required for proper regeneration of adult fins and larval tails. Together, our findings suggest that regeneration is initiated by lineage-specific epigenetic reprogramming, which precedes and enables the activation of downstream regenerative transcriptional programs. This study provides a high-resolution framework for understanding how dynamic chromatin changes govern regenerative plasticity *in vivo* and establishes the EpiTag transgenic line as a powerful new tool for dissecting the epigenetic basis underlying tissue regeneration.

## Results

### The EpiTag epigenetic reporter is an early marker of regeneration

To investigate epigenetic reprogramming during early regeneration, we used a novel *Tg(dazl-ef1a:gfpd2)*^y603^ transgenic zebrafish line our laboratory recently developed that rapidly reports demethylation and other changes in epigenetic marks via altered expression of a destabilized GFP (GFPd2) under the control of the *dazl* CpG island and the Xenopus ef1a promoter (**Fig. 1A**) – hereafter referred to as “EpiTag” fish (Marvel et al., 2025). During normal development this transgene becomes epigenetically silenced by 5 days post fertilization (dpf) and this repression is maintained in all somatic tissues throughout adulthood. Upon caudal fin amputation, however, adult EpiTag zebrafish exhibited transient activation of GFPd2 expression beginning around 16 hours post amputation (hpa) (**Fig. 1B**). This activation peaked between 1 to 2 days post amputation (dpa) and returned to baseline by 10 dpa. Quantification of GFPd2 protein levels using ELISA (**Fig. 1C**), transcript levels using RNA-seq (**Fig. 1D**), and spatial patterns using HCR (**Fig. 1E**) confirmed that GFPd2 proteins peaks at 1 dpa while transcript levels rise earlier, between 8 to 12 hpa. Notably, the well-characterized early regeneration marker *msx1b* (Akimenko et al., 1995; Raya et al., 2003) is co-expressed with EpiTag GFPd2, however *msx1b* expression is initiated later than EpiTag activation (**Fig. 1E**), suggesting epigenetic activation occurs upstream of traditional blastema gene expression. Comparable rapid activation of EpiTag GFPd2 expression is also noted during fin regeneration in 3-7 dpf zebrafish larvae (**Fig. 1F**). EpiTag expression represents the earliest reported marker of cells contributing to fin regeneration, making it an extremely useful “tag” for visualization, identification, and/or isolation of cells at some of the earliest stages of regeneration.

**Figure 1.**
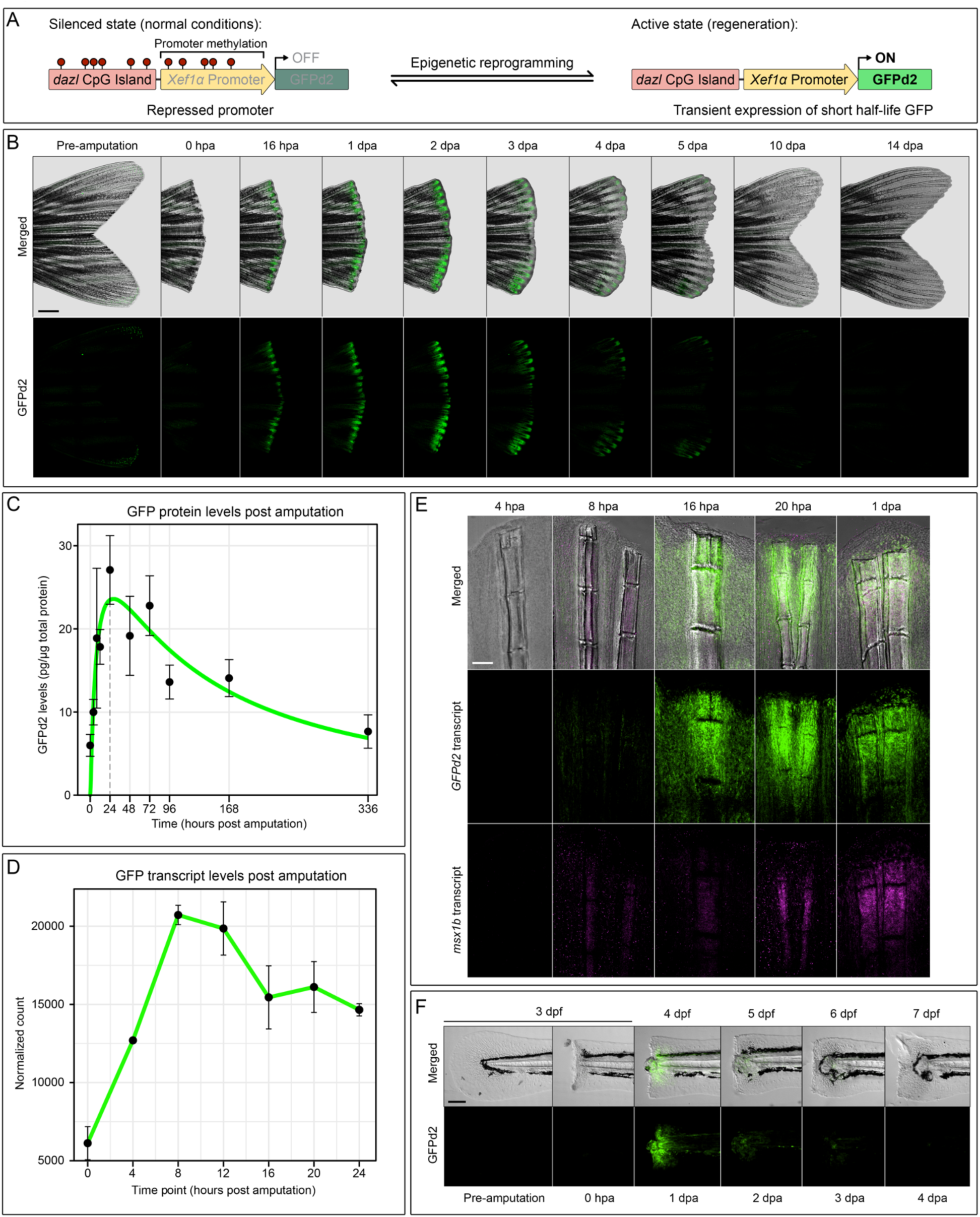
Transient activation of the EpiTag epigenetic reporter during fin regeneration. (A) Schematic representation of the EpiTag transgene construct, consisting of the zebrafish *dazl* CpG island, the Xenopus *ef1a* promoter, and a destabilized GFP (*GFPd2*). Under normal conditions, methylation of the *dazl* CpG island silences promoter activity (OFF). During regeneration, epigenetic reprogramming leads to demethylation and activation of the GFPd2 reporter (ON). (B) Time-lapse fluorescence imaging of an adult EpiTag caudal fin during regeneration, showing basal GFPd2 levels before amputation, followed by transient GFPd2 expression that peaks between 1 to 2 dpa before returning to baseline by 10 dpa. (C) Quantification of GFPd2 protein levels via GFP ELISA across regeneration time points. Protein levels peak at 24 hpa, consistent with reporter imaging. (D) Time-course RNA-seq analysis of GFPd2 transcription during regeneration. Transcription peaks between 8 to 12 hpa, preceding peak protein levels. (E) HCR staining of *GFPd2* (green) and *msx1b* (magenta) transcripts in adult EpiTag fins during the first 24 hours of regeneration. (F) Time-lapse imaging of larval tail regeneration in EpiTag fish from 3 (0 hpa) to 7 dpf (4 dpa).

### Multi-omic analysis of fin regeneration using EpiTag transgenic zebrafish

We used EpiTag fish to carry out a multi-omic analysis of zebrafish fin regeneration, as described in detail in **Fig. 2A** and **Fig S1**. Tissue was excised from amputated fin margins at either 0 hpa or 24 hpa and dissociated into single cells. The 24 hpa cells were FACS sorted to yield 24 hpa GFP+ and 24 hpa GFP-populations, while the 0 hpa total cells were all collected after being run through the FACS machine using the same sorting conditions employed for the 24 hpa cells (**Fig. 2A**). Each of these three cell samples was split and used for matched bulk RNA-seq, ATAC-seq, and bisulfite-seq assays to eliminate the confounding effects of batch or stage variability. We also used additional, separate samples prepared from EpiTag fish to carry out single-cell RNA-seq on 24 hpa GFP+, 24 hpa GFP-, and 0 hpa total dissociated cells, as well as time-course bulk RNA-seq assays on 0, 4, 8, 12, 16, 20, and 24 hpa excised fin margins. Together, this comprehensive set of interlocking omics assays provides a powerful framework for dissecting the temporal and cell-type-specific regulation of regeneration. In this study, we provide examples of how this dataset can be queried to reveal novel biological insights. More importantly, we anticipate that this resource will serve as a foundational tool for the regeneration community to pursue many other questions.

**Figure 2.**
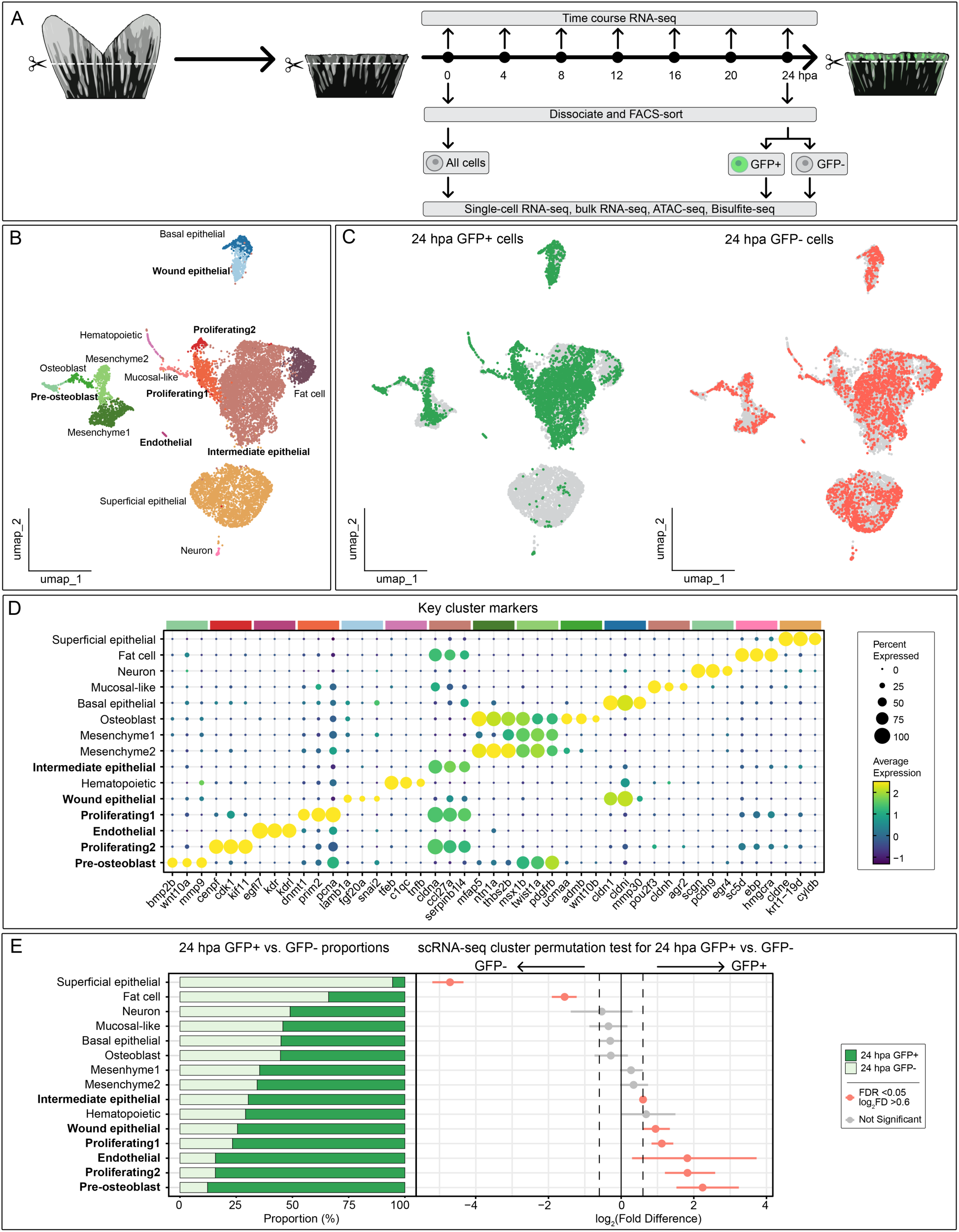
Experimental design and single-cell transcriptomic analysis of regenerating fin tissues. (A) Schematic overview of the multi-omics workflow. Adult EpiTag fins were amputated, and tissues were collected at multiple time points for time-course RNA-seq. At two key time points (0 and 24 hpa), fins were dissociated, FACS-sorted, and used for scRNA-seq, bulk RNA-seq, ATAC-seq, and bisulfite-seq. (B) Integrated UMAP plot of the 24 hpa GFP+ and GFP-cells. (C) UMAP plots split by sample identity reveal differences in the cellular contributions of GFP+, GFP-, and 0 hpa populations to the overall clustering. (D) Dot plot showing representative marker genes used to annotate the major cell types. (E) Bar graph showing the proportion of GFP+ and GFP-cells in each cluster (left). Permutation test results from scProportionTest showing clusters with significant enrichment of GFP+ or GFP-cells (right). Cell clusters noted in bold text in panels D and E show significant enrichment in GFP+ cells.

### Single-cell analysis defines cellular heterogeneity and identifies epigenetically active GFP+ enriched populations

To characterize the transcriptional landscape of early regeneration at single-cell resolution, we performed scRNA-seq on dissociated, FACS-isolated 0 hpa control cells, 24 hpa GFP+ cells, or 24 hpa GFP-cells, each isolated from the caudal margin of regenerating EpiTag fins (**Fig. 2A, Fig. S1**). Integrative UMAP analysis revealed a diverse array of cell types, including superficial, intermediate, and basal epithelial cells, mesenchymal cells, endothelial cells, proliferating cells, hematopoietic populations, mucosal-like cells, fat cells, osteoblasts, pre-osteoblasts, and neurons (**Fig. 2B**). Consistent with previous reports suggesting these cells do not contribute to regeneration (Hou et al., 2020), the superficial epithelial cells were notably depleted from the GFP+ population (**Fig. 2C**). Canonical markers were used to annotate clusters and validate cell identity (**Fig. 2D**). Quantification of GFP+ proportions across clusters showed significant enrichment in regeneration-associated lineages, including pre-osteoblasts, proliferating cells, endothelial cells, wound epithelium, and intermediate epithelium (**Fig. 2E**), indicating that epigenetic activation is confined to a defined subset of regenerative cell types.

Among these, the osteoblast lineage is a particularly well-characterized cell type in fin regeneration. Osteoblasts are known to dedifferentiate into stem-cell-like pre-osteoblasts that contribute to the blastema (Blum & Begemann, 2015; Knopf et al., 2011; I. Sehring et al., 2022). To confirm this transition in our dataset, we performed RNA trajectory analysis of the mesenchymal/skeletogenic partition, which showed that pre-osteoblasts occupy the terminal position along the trajectory, immediately downstream of osteoblasts, which supports their dedifferentiation origin (**Fig. S2A,B**). To further characterize this population, we mined our scRNA-seq data for genes with highly enriched expression in pre-osteoblasts and identified two candidate markers, *abcc6a* and *anos1b* (**Fig. S2C,D**). Whole mount *in situ* hybridization (WMISH) performed on partially amputated fins revealed that *abcc6a* is normally expressed in joint regions of uninjured fins but becomes upregulated in the blastema during regeneration (**Fig. S2E**). In contrast, *anos1b* was undetected in uninjured fins but was also robustly expressed in the regenerating blastema (**Fig. S2F**). Together, these findings suggest that isolation of EpiTag-positive cells from the regenerating fin margin strongly enriches for cell populations actively undergoing regenerative reprogramming.

### RNA-seq reveals temporally distinct gene expression modules in GFP+ cells

To carry out a deeper analysis of genes whose expression was increased in GFP+ cells, we performed bulk RNA-seq on dissociated, FACS-isolated 0 hpa control, 24 hpa GFP+, or 24 hpa GFP-cells, each isolated from the caudal margin of regenerating EpiTag fins (**Fig. 3A, S1, S3**). Differential expression analysis identified 2,489 differentially expressed genes (DEGs) that were significantly increased in both the 24 hpa GFP+ vs. 24 hpa GFP-RNA-seq sample comparison, and in the 24 hpa GFP+ vs. 0 hpa total cell RNA-seq sample comparison. Gene ontology (GO) term analysis of these overlapping DEGs revealed significant enrichment for cell cycle, chromatin organization, and methyltransferase activity (**Fig. 3B**). To broadly assess the temporal dynamics of the 2,489 genes noted above during early regeneration, we examined the expression levels of each of these same genes across a separately generated bulk RNA-seq time-course dataset that was prepared from total, unsorted cells collected from regenerating caudal fin margins at 0, 4, 8, 12, 16, 20, and 24 hpa (**Fig. 3A,C, S1**). Gene expression patterns in this dataset were broadly clustered into four temporally distinct gene expression modules (**Fig. 3D**). Early gene modules (Modules 1 and 2) were enriched for RNA processing, epigenetic regulation, and mitochondrial function, while late-expressing modules (Modules 3 and 4) were associated with cell proliferation and morphogenesis (**Fig. 3E**). These data suggest that epigenetic priming precedes morphogenetic gene expression during early regeneration.

**Figure 3.**
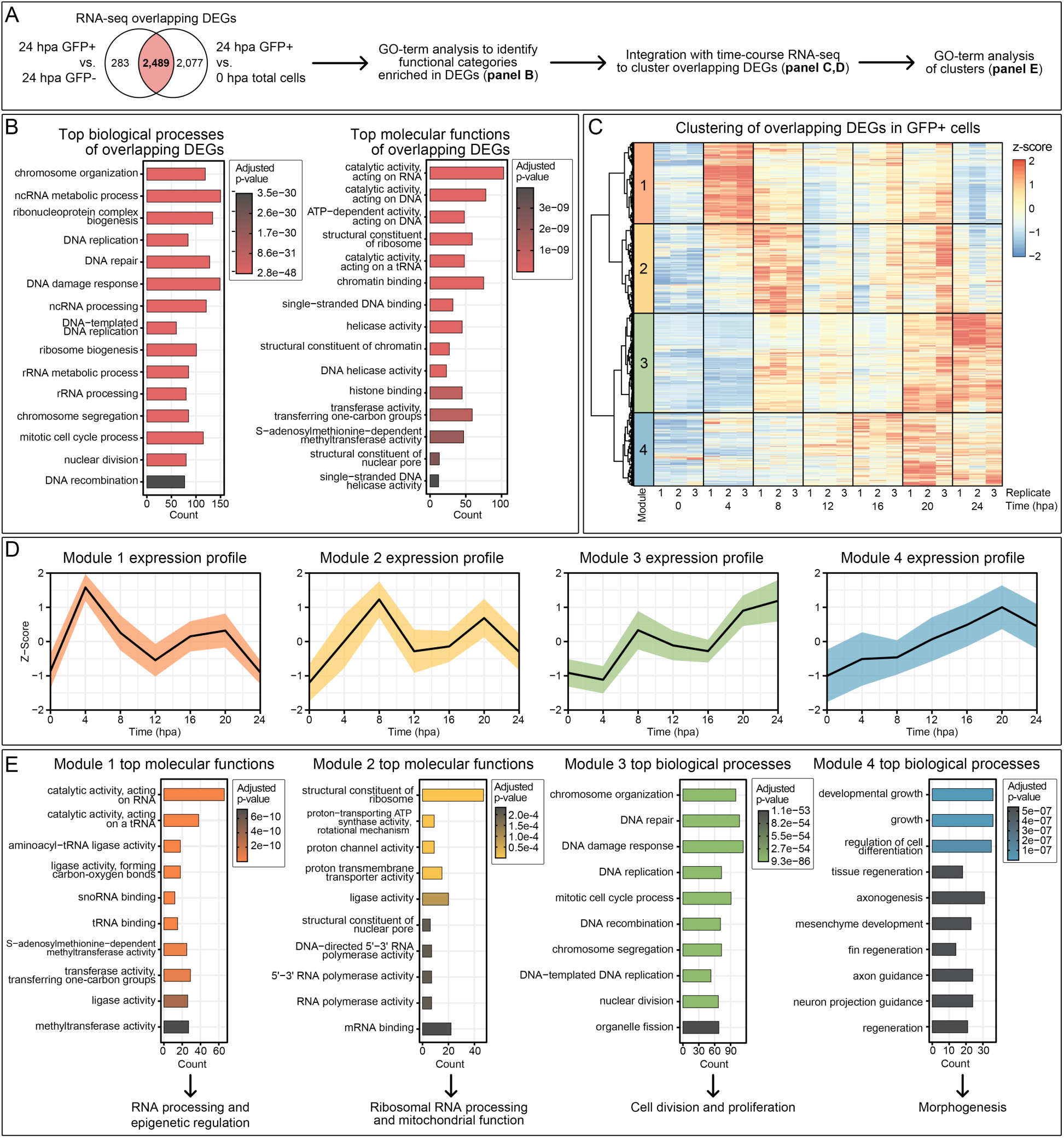
Integrative analysis of bulk and time-course RNA-seq data reveals distinct temporal clusters of genes activated in GFP+ cells. (A) Summary of RNA-seq analysis workflow. Venn diagram illustrates the overlap between differentially expressed genes (DEGs) from GFP+ vs. GFP-and GFP+ vs. 0 hpa control comparisons. (B) GO term enrichment analysis of overlapping DEGs. Bar plots showing the top-enriched biological processes (left) and molecular functions (right). (C) Heatmap of standardized expression profiles (z-scores) for overlapping DEGs across time points. Genes were clustered into four distinct expression modules. (D) Line plots showing the average expression profiles for each of the regeneration gene modules, with shaded regions indicating standard deviation. (E) GO term enrichment of the gene modules.

### Chromatin accessibility changes are associated with morphogenesis genes

To understand changes in chromatin architecture during regeneration, we performed ATAC-seq on the same dissociated, FACS-isolated 0 hpa control, 24 hpa GFP+, or 24 hpa GFP-cell samples that were used for the bulk RNA-seq expression analysis described above (**Fig. 2A, S1**). Differentially accessible regions (DARs) were identified for both 24 hpa GFP+ vs. 24 hpa GFP-and for 24 hpa GFP+ vs. 0 hpa total cell comparisons and were subsequently linked to nearby genes (**Fig. 4A**). For both comparisons, non-coding regions represent the majority (>90%) of DARs (**Fig. S4A**). Genome browser tracks at the *fgf3* and *fgf4* loci revealed changes in chromatin accessibility during regeneration, highlighting regulatory remodeling near these known regenerative genes. In contrast, tracks at the *cldne* locus, a gene highly expressed in superficial epithelial cells but absent from the GFP+ positive population, showed no changes in chromatin accessibility (**Fig. 4B**). A total of 2,714 overlapping DAR-associated genes were also identified from the two comparisons. GO term analysis of these overlapping genes revealed enrichment for functions related to morphogenesis, neurogenesis, and skeletal development (**Fig. 4C**). Transcription factor (TF) footprinting analysis performed using TOBIAS identified candidate TFs with increased binding activity in GFP+ cells (**Fig. S5A**), including known regulators of limb development involved in chromatin remodeling and hedgehog signaling pathways (**Fig. S5B**). Correlating DAR-associated genes with the temporal DEG expression modules identified in the bulk time-course RNA-seq described above (**Fig. 3D**) revealed that “late” module 4 showed a unique and highly significant enrichment for genes associated with DARs (∼8-fold; p<0.0001) compared to the other earlier expression modules (**Fig. 4D**). These data highlight a strong link between chromatin accessibility and the late-appearing morphogenetic programs enriched in module 4, suggesting these morphogenetic programs may be the targets of epigenetic changes initiated during earlier stages of the regenerative process.

**Figure 4.**
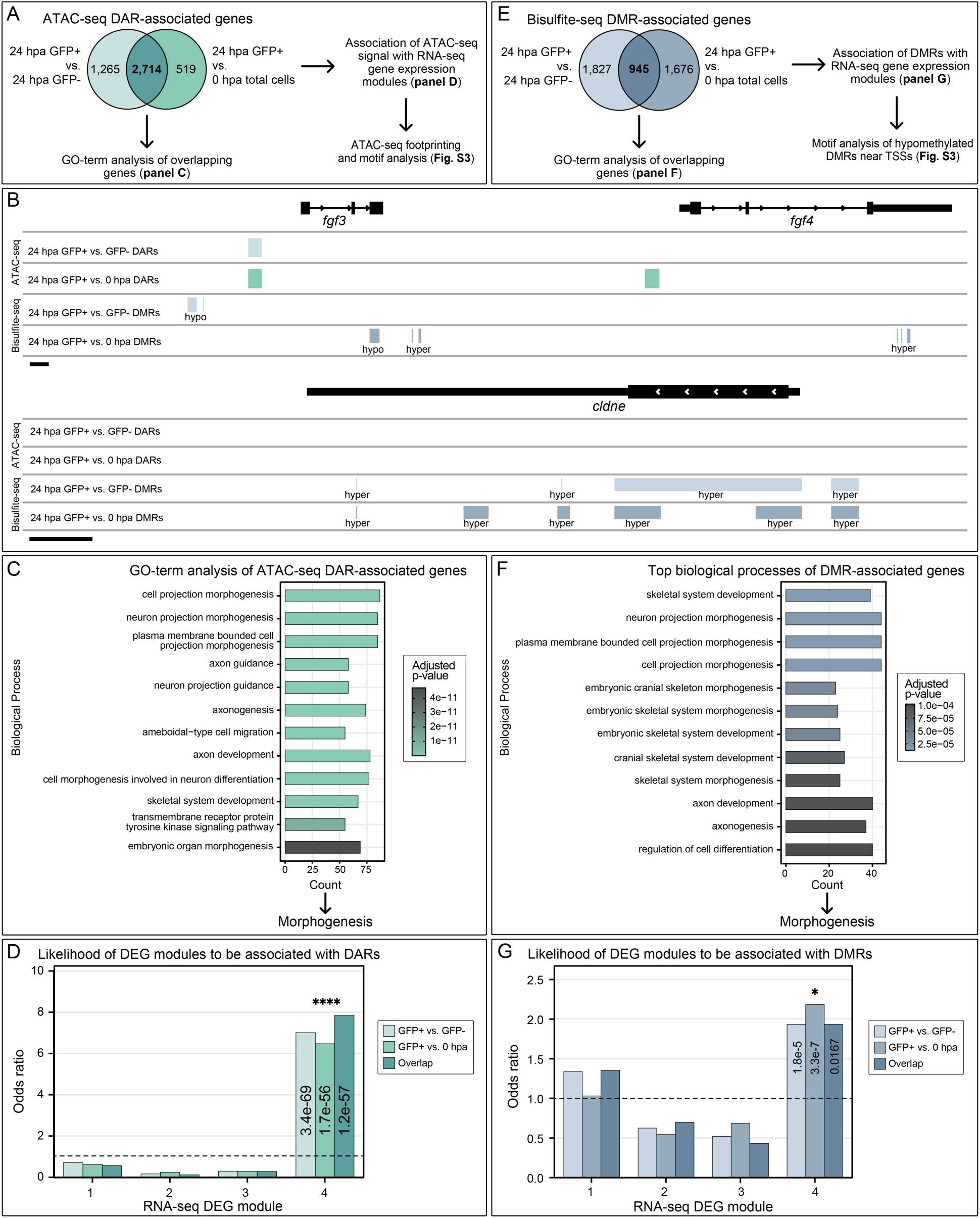
ATAC-seq and bisulfite-seq analysis identifies chromatin accessibility and DNA methylation changes in GFP+ cells. (A) Summary of ATAC-seq analysis workflow. Venn diagram shows genes associated with differentially accessible regions (DARs) from GFP+ vs. GFP-and GFP+ vs 0 hpa comparisons. A total of 2,714 overlapping DAR-associated genes were identified and used for downstream analyses. (B) Genome tracks of the *fgf3* and *fgf4* loci, genes known to be upregulated in the blastema showing increased chromatin accessibility (ATAC-seq) and DNA hypomethylation (bisulfite-seq) during regeneration. In contrast, tracks at the *cldne* locus, a gene expressed in the superficial epithelium (not labeled by EpiTag), show DNA hypermethylation and no DARs in regulatory regions. (C) GO term enrichment of the overlapping DAR-associated genes, showing enrichment for processes related to tissue morphogenesis and development. (D) Chi-squared analysis of DAR association for each of the four DEG clusters identified in Figure 3 (****p<0.0001). (E) Summary of bisulfite-seq analysis workflow. Venn diagram shows differentially methylated regions (DMRs) identified from 24 hpa GFP+ vs. GFP- and 24 hpa GFP+ vs 0 hpa comparisons. A total of 945 overlapping DMR-associated genes were identified and used for downstream analyses. (F) GO term enrichment of the overlapping DMR- associated genes, showing enrichment for processes related to morphogenesis and skeletal development. (G) Chi-squared analysis of DMR association for each of the four DEG clusters identified in Figure 3 (*p<0.05).

### DNA methylation changes are also associated with morphogenesis genes

To explore the role of DNA methylation during regeneration, we performed bisulfite sequencing on the same dissociated, FACS-isolated 0 hpa control, 24 hpa GFP+, or 24 hpa GFP- cell samples that were used for the bulk RNA-seq and ATAC-seq analyses described above. Differentially methylated regions (DMRs) were identified for both 24 hpa GFP+ vs. 24 hpa GFP- and for 24 hpa GFP+ vs. 0 hpa total cell comparisons and linked to nearby genes (**Fig. 4E**). For both comparisons, non-coding regions represent the majority (>90%) of DMRs, with approximately 30% distributed around promoter regions (**Fig. S4B**). DNA methylation tracks revealed hypomethylated regions near *fgf3* and *fgf4*, genes upregulated in the blastema. In contrast, the *cldne* locus, which is absent from GFP+ populations, showed hypermethylation across its regulatory regions (**Fig. 4B**). We compared DMR-associated genes in both 24 hpa GFP+ vs. 24 hpa GFP- and 24 hpa GFP+ vs. 0 hpa total cell control datasets and identified a total of 945 shared genes. GO term analysis revealed that these shared genes were enriched for morphogenetic and developmental processes such as skeletogenesis and neurogenesis (**Fig. 4F**). By again correlating DMR-associated genes with the temporal DEG expression modules identified in the bulk time-course RNA-seq described above (Fig. 3C-E), we found that “late” module 4 was also enriched for genes associated with DMRs (∼2 fold; p<0.05) (**Fig. 4G**), consistent with ATAC-seq trends. Additionally, *de novo* motif enrichment analysis of hypomethylated DMRs near TSSs revealed enrichment for candidate TF binding sites with known roles in skeletogenesis, neurogenesis, and limb bud development (**Fig. S5C**), supporting a role for targeted epigenetic remodeling in facilitating activation of regeneration-primed morphogenesis gene expression programs.

### Epigenetic regulators from chromatin remodeling and histone-modifying complexes are activated in the early blastema

Our multi-omics analysis suggests that early epigenetic “priming” via upregulated expression of epigenetic regulators may be important for subsequent expression of regenerative gene expression programs. To identify candidate epigenetic regulators mediating regeneration, we identified epigenetic-related genes highly expressed in 24 hpa GFP+ cells and enriched in early DEG modules (Modules 1 and 2). These included members of the ATP-dependent chromatin remodeling complexes, the Inhibitor of Histone Acetyltransferase (INHAT) complex, and the Facilitates Chromatin Transcription (FACT) complex. Single-cell analysis revealed strong expression of these candidates in proliferating cells and pre-osteoblast clusters (**Fig. 5A**). Bulk RNA-seq analysis confirmed their upregulation in GFP+ cells (**Fig. 5B**). Moreover, whole-mount *in situ* hybridization (WMISH) validated their spatial localization in the early blastema of the adult regenerating fin (**Fig. 5C**). All candidates were also upregulated in the notochord bead during larval tail fin regeneration (**Fig. 5D**). These data suggest that genes with known roles in epigenetic reprogramming are strongly expressed at early stages of fin regeneration, and that the epigenetic function of these genes may be required for activation/regulation of subsequent regenerative processes.

**Figure 5.**
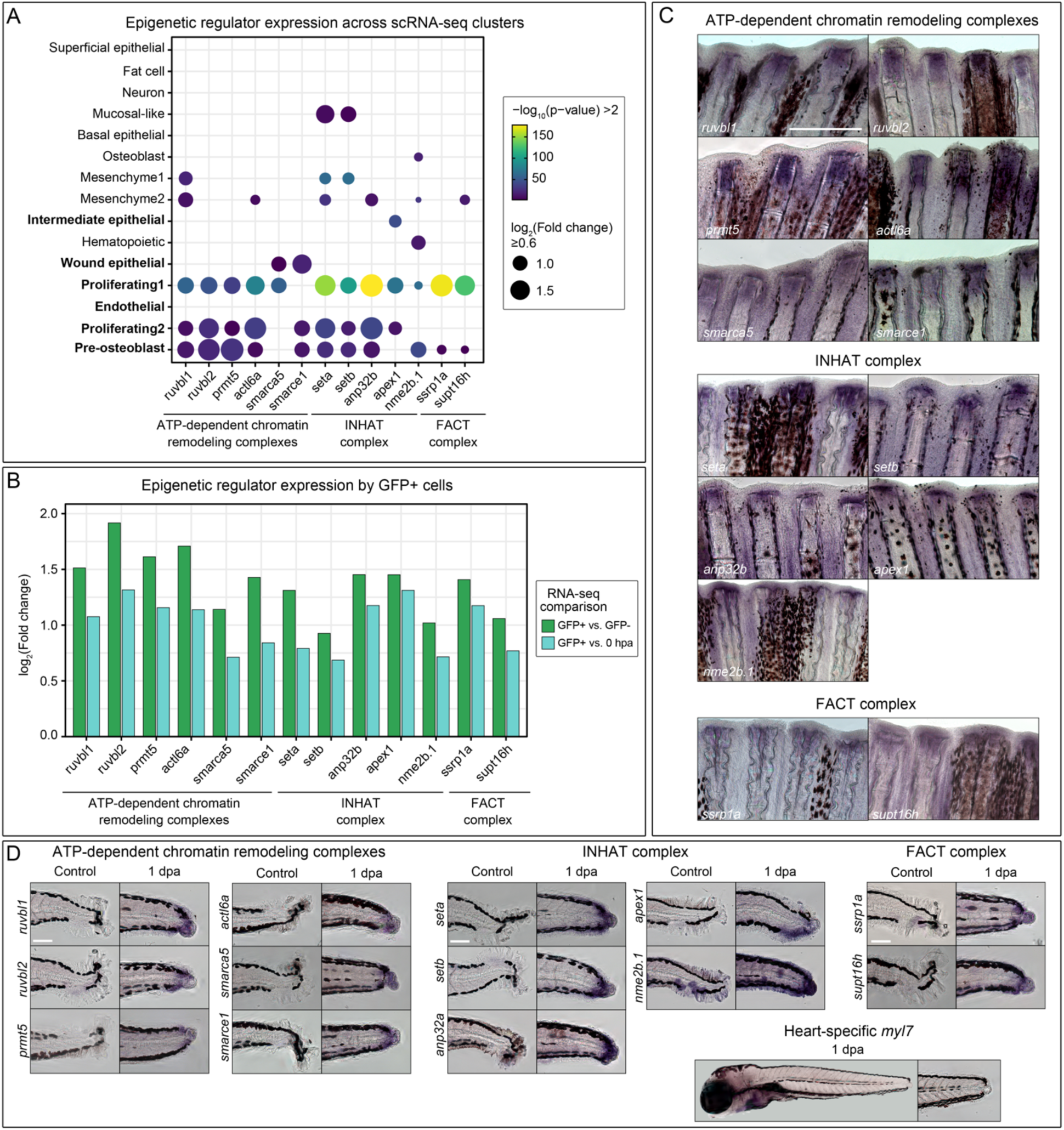
Regeneration-specific epigenetic regulators are upregulated in the early blastema. (A) Dot plot showing the expression of candidate epigenetic regulators from the ATP- dependent chromatin remodeling complexes, the Inhibitor of Histone Acetyltransferase (INHAT) complex, and the Facilitates Chromatin Transcription (FACT) complex across GFP+ scRNA-seq clusters. Cell clusters noted in bold text are those that show significant enrichment in GFP+ cells. (B) Bar graph showing differential expression of the same genes from RNA-seq comparisons (24 hpa GFP+ vs. GFP- and 24 hpa GFP+ vs. 0 hpa). (C) Whole-mount *in situ* hybridization (WMISH) of adult fins at 1 dpa confirms blastema-localized expression of the epigenetic regulators. Scale bar, 0.5 mm. (D) WMISH of larval tail fins at 1 dpa showing similar regeneration-induced expression for all genes except *smarcad1b*. Scale bar, 100 μm. A heart-specific gene (*myl7*) was included as a negative control and shows no expression in the regenerating tail fin.

### Functional inhibition of *ruvbl1* and *ruvbl2* impairs adult caudal fin and larval tail regeneration

In order to test the functional importance of a few of the epigenetic regulators we identified as being upregulated during early regeneration, we focused on *ruvbl1* and *ruvbl2*, key members of multiple ATP-dependent chromatin remodeling complexes. The expression of each of these genes is strongly increased in 24 hpa GFP+ cells (**Fig. 5A**), co-localizes with GFPd2 transcripts in the 1 dpa regenerating fin blastema (**Fig. 6A**), and shows upregulation in the bulk time-course RNA-seq (**Fig. 5B**) To assess the roles of Ruvbl1 and Ruvbl2 in fin regeneration, we performed targeted knockdown using Vivo-Morpholinos (Vivo-MOs) injected into amputated adult fins (**Fig. 6B**). Injection conditions were optimized using a GFP Vivo-MO control, which effectively knocked down EpiTag GFP expression in the regenerating fin blastema for up to three days after a single injection at the caudal fin margin at 4 hpa (**Fig. S6**). This validated the effectiveness and stability of the Vivo-MO injection for short-term gene knockdown. Knockdown of either *ruvbl1* or *ruvbl2* significantly reduced regenerative growth compared to control Vivo-MO-injected fins by at least 20% (**Fig. 6C,D**). To complement this approach, we also performed pharmacological inhibition of the Ruvbl1/2 complex in regenerating larval tails using the small molecule inhibitor CB6644, a selective allosteric small-molecular inhibitor of the ATPase activity of the Ruvbl1/2 complex with anticancer activity (Assimon et al., 2019; Yi et al., 2024) (**Fig. 6E**). CB6644 treatment impaired regeneration in a dose-dependent manner (**Fig. 6F,G**). Together, these results suggest that Ruvbl1/2 activity is required for proper regenerative growth in both adult and larval contexts, likely via its function in ATP-dependent chromatin remodeling.

**Figure 6.**
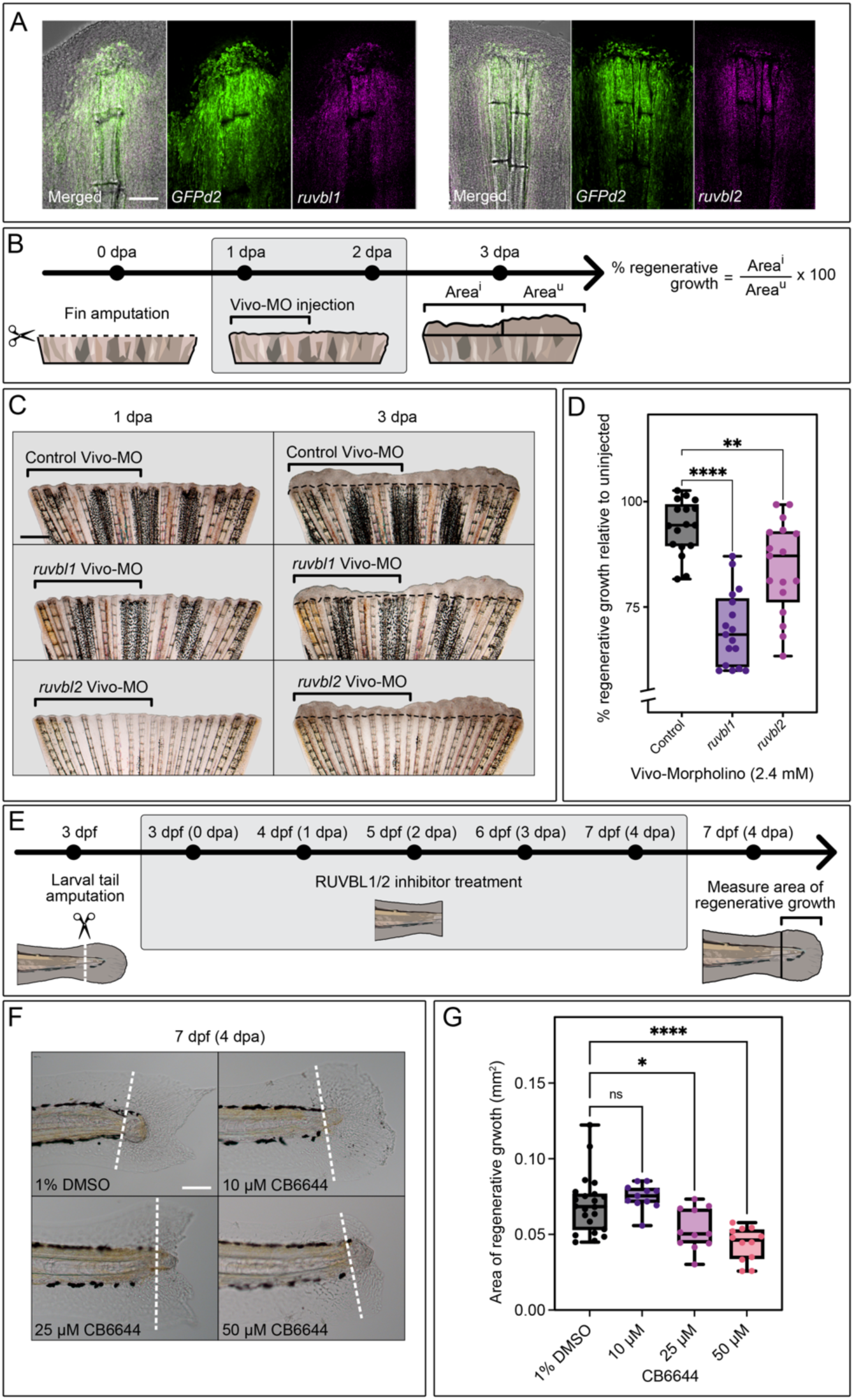
Functional validation of ruvbl1/2 activity during regeneration. (A) Hybridization chain reaction (HCR) imaging shows co-localization of *ruvbl1* and *ruvbl2* transcripts (magenta) with *GFPd2* (green). Scale bar, 100 μm. (B) Schematic representation of the Vivo-Morpholino (Vivo-MO) injection strategy in regenerating adult caudal fins. Regenerative growth was quantified at 3 dpa by comparing the injected region (Area^I^) to the uninjected control region (Area^U^) within the same fin. (C) Representative images of adult fins at 1 dpa (pre-injection) and at 3 dpa following injection with control, *ruvbl1*, or *ruvbl2* Vivo-MOs at 1 and 2 dpa. Knockdown of *ruvbl1* or *ruvbl2* resulted in reduced regenerative outgrowth. Scale bar, 0.5 mm. (D) Violin plots showing percent regenerative growth across different conditions (****p<0.0001; **p<0.01). (E) Schematic representation of larval tail regeneration inhibitor treatment strategy. Tails were amputated at 3 dpf and regeneration was monitored in the presence of Ruvbl1/2 inhibitor CB6644 from 0 to 4 dpa (3 to 7 dpf). (F) Representative images of regenerating larval tails treated with 1% DMSO (vehicle control) or CB6644 at 10, 25, or 50 μM. Scale bar, 100 μm. (G) Violin plots showing regenerative area (mm^2^) across treatments (ns, not significant; *p<0.05; ****p<0.0001).

## Discussion

Understanding how epigenetic reprogramming contributes to tissue regeneration remains a major challenge, in part due to the lack of tools to visualize dynamic and often transient epigenetic changes *in vivo*. To address this, we leveraged the novel EpiTag zebrafish line, a transgenic reporter that uses the *dazl* CpG island and the ubiquitous *Xenopus ef1*α promoter to drive expression of a short-lived GFP (GFPd2) in response to epigenetic changes (Marvel et al., 2025). During normal development, this transgene is rapidly silenced by DNA methylation and GFPd2 expression is undetectable in somatic tissues from 5 days post fertilization (dpf) to adulthood. Upon caudal fin amputation, however, we observed robust and transient activation of the EpiTag reporter beginning at 16 hours post amputation (hpa), peaking between 1 to 2 dpa before returning to baseline expression levels by 10 dpa (**Fig. 1B**). This defined temporal window captures early epigenetic reprogramming well before the expression of canonical blastema markers.

To investigate the molecular features of early epigenetic activation, we combined the EpiTag reporter with a multi-omics approach. FACS-sorted 24 hpa GFP+, 24 hpa GFP- cells, and 0 hpa total cells were profiled by bulk RNA-seq, ATAC-seq, and bisulfite-seq. In parallel, time-course RNA-seq and single-cell RNA-seq were performed to capture dynamic and cell-type-specific transcriptional changes. This integrative strategy enabled high-resolution analysis of chromatin accessibility, methylation levels, and gene expression dynamics from identical regenerative cell populations. Our findings suggest that epigenetic reprogramming is not a downstream consequence of regeneration but is among the earliest events triggered in response to injury. Indeed, GFPd2 transcript and protein levels peaked before *msx1b* expression (**Fig. 1E**), a key transcription factor widely used as a marker of the regenerative blastema in zebrafish fin and heart regeneration (Akimenko et al., 1995; Raya et al., 2003). Notably, orthologs of *msx* genes are also activated during regeneration in axolotl (Carlson et al., 1998), Xenopus (Barker & Beck, 2009), and mice (Taghiyar et al., 2017), pointing to a conserved role in regeneration across species.

A previous study by Lee et al. (2020) profiled DNA methylation during zebrafish fin regeneration but concluded that methylation states are largely maintained in a lineage-specific manner during regeneration. However, their analysis was performed at 4 dpa, likely past the critical window for detecting early epigenetic changes. Moreover, they focused on osteoblasts isolated using a *sp7* transgenic line, which is a terminal differentiation marker that is known to be downregulated during dedifferentiation (Stewart et al., 2014). By using the EpiTag reporter, we were able to capture epigenetically active cells at an earlier stage and across diverse lineages, providing us with a unique window into the first epigenetic changes that set the stage for later events during regeneration. Single-cell RNA-seq identified major cell types consistent with previously published findings on the cellular diversity of the regenerating fin (Hou et al., 2020). We found that EpiTag activation is highly restricted to specific regenerative lineages, including pre-osteoblasts, proliferative cells, endothelial cells, wound epithelium, and intermediate epithelium. Despite being transcriptionally active, superficial epithelial cells showed little to no GFPd2 activation, supporting the view that epigenetic reprogramming is not a broad stress response but a targeted event occurring in regeneration-competent cell populations.

To explore the transcriptional landscape associated with EpiTag activation, we performed bulk RNA-seq on GFP+ and GFP- cells and integrated this with a time-course RNA-seq dataset spanning 0 to 24 hpa. We identified a core set of 2,489 genes that were consistently upregulated in GFP+ cells. Temporal clustering of these genes uncovered four distinct expression modules. Remarkably, early expression modules (Modules 1 and 2) were enriched for genes associated with RNA processing, methyltransferase activity, and mitochondrial function, while late expression modules (Modules 3 and 4) were involved in proliferation and morphogenesis. These findings suggest a regulatory cascade in which early activation of epigenetic remodeling primes cells for subsequent activation of developmental and regeneration gene programs.

Chromatin accessibility and DNA methylation changes mirrored these transcriptional dynamics. ATAC-seq and bisulfite-seq analysis identified differentially accessible regions (DARs) and differentially methylated regions (DMRs) in GFP+ cells, respectively. DARs and DMRs were also found to be enriched near genes related to neurogenesis, skeletogenesis, and morphogenesis. Transcription factor footprinting and motif analyses of these regions identified candidates such as *fos*, *jun*, *zic*, *twist*, *runx*, *gli*, and *runx* family genes, which have established roles in development and regeneration (Chen et al., 2025; Jiang et al., 2021). For example, *gli* and *zic* transcription factors have been reported to regulate skeletogenesis by controlling hedgehog signaling and *runx2* expression (Shimoyama et al., 2007). Additionally, genes in the late expression module (Module 4) were significantly more likely to be associated with both DARs and DMRs, suggesting that early epigenetic regulators may prime these loci for later transcriptional activation.

Within the early expression gene modules, we identified several epigenetic regulators upregulated in GFP+ cells and spatially localized to the blastema. These included components of ATP-dependent chromatin remodeling complexes (e.g. *ruvbl1, ruvbl2*), the INHAT complex (e.g. *seta, setb*), and the FACT complex (e.g. *ssrp1a*). WMISH confirmed their localization to regenerative tissues and co-localization with GFPd2 was observed for *ruvbl1* and *ruvbl2* using HCR. These two factors are evolutionarily conserved AAA+ ATPases that form a hetero-hexameric complex (Gorynia et al., 2011) and are core components of many chromatin remodeling complexes, including INO80, SWR1, TIP60, and p400 (Dauden et al., 2021). Morpholino-mediated knockdown of *ruvbl1* and *ruvbl2* during early development disrupted heart growth in zebrafish larvae (Rottbauer et al., 2002).

To test the functional requirement of these regulators, we performed loss-of-function studies in both adult and larval zebrafish. Vivo-Morpholino (Vivo-MO) knockdown of *ruvbl1* and *ruvbl2* following fin amputation significantly impaired regenerative growth. Similarly, pharmacological inhibition of the Ruvbl1/2 complex using the selective inhibitor CB6644 also reduced regeneration in larval tails in a dose-dependent manner. CB6644 has been previously shown to alter chromatin accessibility and impair proliferation in cancer models (Assimon et al., 2019; Yi et al., 2024). Our findings establish a critical role for Ruvbl1/2 in driving early regeneration, likely through their functions in chromatin remodeling and epigenetic reprogramming.

Taken together, our data support a model in which regeneration is initiated by a transient but tightly regulated wave of epigenetic reprogramming that precedes the transcription of classical regeneration markers. These early changes include coordinated remodeling of chromatin accessibility and DNA methylation that prime key regenerative genes for activation. Unlike development, which relies on progressive and largely irreversible epigenetic states (Bestor et al., 2015), regeneration requires dynamic epigenetic reprogramming to reactivate silenced developmental programs. For example, Duong et al. (2024) reported that embryonic histone methylation patterns are recapitulated during fin regeneration, highlighting the reactivation of developmental gene regulatory networks. The timing and specificity of these epigenetic changes may represent key determinants of regenerative potential in other tissues and organisms.

By integrating a novel epigenetic reporter with multi-omics profiling and functional perturbation, our study provides a comprehensive view of early regenerative reprogramming. Moving forward, dissecting how Ruvbl1/2 and other early epigenetic complexes coordinate signaling pathways and transcription factor expression will be key to understanding how regenerative competence is established. Importantly, because these factors act upstream in the regulatory hierarchy, targeting them could offer therapeutic strategies to potentially reactivate entire networks of tissue repair genes and unlock latent regenerative potential in mammals.

## Materials and Methods

### Zebrafish husbandry and procedures

The zebrafish strain used in this study was *Tg(dazl-ef1a: gfpd2)^y603^* (detailed characterization in Marvel et al., 2025). Adult zebrafish (6 to 12 months old) were anaesthetized in 0.1% tricaine and caudal fins were amputated with a scalpel at approximately 50% of the fin length. Larval tail amputations were performed as previously described (Scott et al., 2022). Briefly, 3 to 5 dpf larvae were anaesthetized in 0.1% tricaine and tail amputations were performed distal to the caudal vein circulatory loop, using the pigment gap as a visual guide. Following amputation, adult and larval zebrafish were allowed to regenerate at 28.5°C. Expression of GFPd2 in regenerating fins was imaged on a Nikon Ti2 inverted microscope with Yokogawa CSU-W1 spinning disk confocal (Hamamatsu Orca Fusion-BT camera). All animal procedures were conducted in an AAALAC-accredited facility under the oversight of the NICHD Animal Care and Use Committee (Animal Study Proposal #24-015).

### Biochemical assays

GFP protein levels were quantified using the Abcam GFP ELISA Kit (cat. no. ab171581). Briefly, amputated fins were collected at various time points and total protein was extracted using the lysis buffer from the kit, supplemented with Halt™ Protease Inhibitor Cocktail, EDTA-Free (Thermo Scientific, cat. no. 78425). Protein concentration was then determined using the Pierce™ BCA Protein Assay Kit (Thermo Scientific, cat. no. 23225). For ELISA, 25 μg of total protein per sample was used, following the manufacturer’s protocol. Optical density (O.D.) of the colorimetric reaction was then measured at 450 nm using a Synergy H1 Microplate Reader (Biotek).

### Tissue dissociation, cell enrichment, and multi-omics workflow

For each biological replicate, regenerating fins (20 fins at 1 dpa) and control fins (3 fins at 0 hpa) were collected in separate 1.5 mL Eppendorf tubes containing 1 mL of 1X Dulbecco’s PBS (DPBS) (Gibco, cat. no. 14190144). Fins were pelleted by centrifugation, washed once with 1X DPBS, and minced directly in the tube using micro-dissecting scissors. The minced tissue was pelleted again and resuspended in 1.4 mL of Accumax Dissociation Solution (Innovative Cell Technologies Inc., cat. no. AM105). Tissue was dissociated by pipetting with a 1000 μL pipette for up to an hour, with periodic checks under the microscope. To improve cell dissociation, the suspension was allowed to settle at the 20-minute mark and the supernatant was transferred to new tubes and topped off with fresh Accumax to dilute the tissue load and facilitate more efficient dissociation.

The remaining bone fragments and undigested tissue were also resuspended in fresh Accumax and incubated further to promote more complete dissociation.

Once dissociation was complete, the samples were pelleted at 700 g for 5 minutes, washed in filter-sterilized wash buffer [1X DPBS supplemented with 2% fetal bovine serum (FBS) and 1% bovine serum albumin (BSA)], and centrifuged again. Cell pellets were pooled into filter-sterilized cell medium [FluoroBrite DMEM (ThermoFisher, cat. no. A1896701) supplemented with 10% FBS and 1% BSA]. Samples containing bone fragments were filtered through a 20 μm cell strainer, while samples containing mostly cells were filtered through a 40 μm cell strainer, followed by rinsing with additional cell medium.

Cell viability was determined using a 20 μL aliquot stained with acridine orange/propidium iodide (Logos Biosystems, cat. no. F23011) and quantified using a LUNA-FL™ Dual Fluorescence Cell Counter (Logos Biosystems, cat. no. L20001). Samples with >90% viability were then used for FACS-sorting. Cells were sorted into GFP+ and GFP- populations using a FACS ARIA III (BD Biosciences). Gating parameters were established using dissociated 0 hpa fins, which lack GFP expression. Post-sort viability was re-assessed and samples with >80% viability and ≥1.5X10^5^ cells were selected for downstream DNA and RNA extraction. Total DNA and RNA were isolated using the Qiagen AllPrep® DNA/RNA Micro kit (cat. no. 80284). In parallel, 50,000 live sorted cells were also allocated for ATAC-seq library preparation. This procedure was repeated across four independent experiments to generate four biological replicates.

### Single-cell RNA-seq library preparation and sequencing

Single-cell suspensions for GFP+, GFP- and 0 hpa samples were loaded on the Chromium X Controller (10x Genomics) using the Chromium Single Cell 3’ v3.1 Kit to generate single-cell gel bead-in-emulsions (GEMs). Reverse transcription, cDNA amplification, and library preparation were carried out according to the manufacturer’s protocol. Libraries were quantified using a Qubit Fluorometer and quality-checked on an Agilent Bioanalyzer. Sequencing was performed on an Illumina NovaSeq 6000 platform.

Raw sequencing data were processed using the Cell Ranger pipeline with alignment to the Danio rerio GRCz11 genome (Ensembl release 99) using the Lawson Lab Zebrafish Transcriptome Annotation v4.3.2 (Lawson et al., 2020). Filtered gene-barcode matrices were analyzed in R (v4.3.1) using Seurat v5.0.1. Low-quality cells were removed based on thresholds for gene count (<200 or>6000), mitochondrial content (>10%) and doublet detection using scDblFinder (v1.12.0). Data were log-normalized and scaled and highly variable genes were identified using the “vst” method. Dimensionality reduction was performed using PCA, followed by UMAP for visualization.

For integrated analysis, anchors were identified using FindIntegrationAnchors. Graph-based clustering was performed using FindNeighbors (dims = 1:30) and FindClusters (resolution = 0.3, algorithm = 4). Specific clusters were further subclustered using FindSubCluster at a resolution of 0.15 on the parent clustering graph. Subclusters were visualized using UMAP and used for downstream analysis. Differentially expressed genes (DEGs) for each cluster were identified using FindAllMarkers on the RNA assay. Cluster proportions were compared across conditions using scProportionTest package (v0.0.0.9). RNA trajectory analysis was performed using Monocle3 (v1.3.7).

### RNA-seq library preparation and analysis

RNA quality was assessed using an Agilent BioAnalyzer and samples with RIN >8 were used to generate libraries using the Zymo-Seq RiboFree Total RNA Library Kit (Zymo, cat. no. R3000) and sequenced on an Illumina NovaSeq 6000 platform (2X100 bp paired-end reads). Libraries were quantified using a Qubit Fluorometer and quality-checked on an Agilent Bioanalyzer. Adapter and quality trimming were performed using Trimmomatic (v0.39) and reads were aligned to the Danio rerio GRCz11 genome (Ensembl release 99) using HISAT2 (v2.2.1). Gene counts were obtained using featureCounts (Subread v2.0.3) with the Ensembl GTF annotation (release 99). Differential expression analysis was performed using DESeq2 (v1.40.2). Genes with adjusted p-value <0.05 and log₂(Fold change) ≥0.6 were considered differentially expressed for the comparison between GFP+ and GFP- and genes with adjusted p-value <0.005 and log₂(Fold change) ≥0.6 were considered differentially expressed for the comparison between GFP+ and 0 hpa.

### ATAC-seq library preparation and analysis

ATAC-seq libraries were generated using the Zymo-Seq ATAC Library Kit (Zymo Research, cat. no. D5458). Libraries were quantified using a Qubit Fluorometer and quality-checked on an Agilent Bioanalyzer. Libraries were sequenced on an Illumina NovaSeq 6000 platform (2X50 bp paired-end reads). Reads were trimmed using Trimmomatic (v0.39) and aligned to the Danio rerio GRCz11 genome using Bowtie2 (v2.4.5). PCR duplicates and mitochondrial reads were removed using Picard and samtools. Accessible chromatin peaks were called using MACS2 (v2.2.7.1). Peak count matrices were generated and analyzed with DiffBind (v3.10.1). Peaks were filtered and merged across replicates and sample-specific read counts were normalized using the DESeq2 method. Principal component analysis (PCA) was used for initial quality control.

Differential chromatin accessibility was assessed across GFP+ vs GFP- and GFP+ vs 0 hpa control comparisons. Differentially accessible regions (DARs) were defined as those with FDR <0.1 (GFP+ vs. GFP-) or <0.01 (GFP+ vs. 0 hpa) and log₂(Fold change) ≥0.6. DARs were exported as BED files for downstream analyses. To identify transcription factor (TF) activity changes, TOBIAS (v0.13.3) was used for ATAC-seq footprinting analysis. Signal tracks were corrected for Tn5 insertion bias using ATACorrect and binding scores were calculated with ScoreBigwig. Differential TF binding between conditions was identified using BINDetect and footprinting volcano plots were generated using ggplot2.

### Bisulfite-seq library preparation and analysis

Genomic DNA was bisulfite-converted using the Pico Methyl-Seq™ Library Prep Kit (Zymo Research, cat. no.). Libraries were quantified using a Qubit Fluorometer and quality-checked on an Agilent Bioanalyzer. Sequencing was performed on an Illumina NovaSeq 6000 platform (2X50 bp paired-end reads). Adapter trimming was performed with Trimmomatic (v0.32) and reads were aligned to a bisulfite-converted Danio rerio GRCz11 reference genome using Bismark (v0.23.1). Duplicate reads were removed and methylation calls were extracted using Bismark Methylation Extractor, generating per-base CpG methylation reports for each sample.

CpG methylation data were imported into R using bsseq (v1.36.0). Smoothed methylation profiles were generated using BSmooth with default parameters. For each comparison (GFP+ vs GFP- and GFP+ vs 0 hpa control), CpG loci were filtered to retain sites with ≥3X coverage in at least three replicates per group. Differentially methylated regions (DMRs) were identified using BSmooth.tstat, followed by dmrFinder, with thresholds set to the top 2.5% of the t-statistic distribution, requiring DMRs to contain ≥5 (GFP+ vs. GFP-) or ≥8 (GFP+ vs. 0 hpa) CpGs with an absolute mean methylation difference ≥0.1.

For motif analysis, hypomethylated DMRs located near transcription start sites (+3 kb) were selected and further filtered by coverage. In the GFP+ vs GFP- comparison, DMRs with ≥5X coverage were used, while for GFP+ vs 0 hpa, DMRs with ≥8X coverage were included. Enriched sequence motifs within these regions were identified using the STREME algorithm from the MEME Suite (v5.5.5), and motif similarity searches were performed using Tomtom against known transcription factor databases to identify candidate regulators.

### Time-course RNA-seq library preparation and analysis

Caudal fin tissues were collected at seven time points following amputation (0, 4, 8, 12, 16, 20, 24 hpa). For each time point, 8 fins were collected for each of the four biological replicates. Total RNA was extracted using the Qiagen RNeasy Micro Kit (cat. no. 74004) and libraries were prepared using the Zymo-Seq RiboFree Total RNA Library Kit (Zymo Research, cat. no. R3000) following the manufacturer’s protocol. Libraries were sequenced on an Illumina NovaSeq 6000 platform (2X100 bp paired-end reads). Raw reads were trimmed using Trimmomatic (v0.39) and aligned to the Danio rerio GRCz11 reference genome (Ensembl release 99) using HISAT2 (v2.2.1). Gene-level counts were obtained using featureCounts (Subread v2.0.3) with the Ensembl GTF annotation (release 99). Normalization was performed using DESeq2 (v1.40.2) and genes previously identified as differentially expressed in bulk RNA-seq comparisons (GFP+ vs GFP- and GFP+ vs 0 hpa) were selected for downstream analysis. Gene Ontology enrichment analysis for selected clusters was performed using clusterProfiler (v4.6.0) with the org.Dr.eg.db annotation package. The proportion of DEGs overlapping with ATAC-seq peaks was evaluated using Chi-squared tests with Benjamini-Hochberg correction.

### Whole mount *in situ* hybridization and hybridization chain reaction

Whole mount *in situ* hybridization (WMISH) was performed as previously described with minor modifications (Marvel et al., 2025). Briefly, amputated fins were fixed in 4% paraformaldehyde (PFA) over 2 days at 4°C. Adult fins were digested with 10 μg/mL proteinase K for 1 hour, while 5 dpf larvae were digested for 30 minutes. WMISH samples were imaged using a Leica DMI 8000B inverted compound microscope with Leica DMC6200 camera. Hybridization chain reaction (HCR) was also performed as previously described with minor modifications (Marvel et al., 2025). Following fixation and overnight methanol incubation, amputated fins were bleached in 0.18 mM KOH in methanol for 5 minutes. After rehydration, the fins were digested with 10 μg/mL proteinase K for 20 minutes. HCR probes targeting *GFPd2*, *msx1b*, *ruvbl1*, and *ruvbl2* were designed in-house and synthesized as 50 pmole oligo pools (oPool) from Integrated DNA Technologies (Coralville, IO, USA). HCR samples were imaged using the previously described spinning disk confocal microscope.

### Vivo-Morpholino knockdown

Antisense Vivo-Morpholinos (Vivo-MOs) (Gene Tools) were reconstituted at a stock concentration of 4 mM. Injections were carried out using a final concentration of 2.4 mM Vivo-MO mixed with 1X PBS and 0.1% Texas Red™ Dextran (10,000 MW) (Invitrogen, cat. no. D1828). The translation-blocking Vivo-MO sequences were: ruvbl1 5’-ACTTCTTCGATCTTCATGTTTCTTA-3’ (Huang et al., 2013) and ruvbl2 5’-TTGCCACCTGCGCTGCCATGTTTTC-3’ (Zhao et al., 2013). A standard control Vivo-MO: 5’-CCTCTTACCTCAGTTACAATTTATA-3’ and a Vivo-MO targeting GFP: 5’-ACAGCTCCTCGCCCTTGCTCACCAT-3’ from Gene Tools were used as negative and positive controls, respectively. Injections were performed using a Pneumatic PicoPump (World Precision Instruments) into the dorsal side of regenerating caudal fins at 1 and 2 dpa. The ventral side of the fins was left uninjected and served as internal controls. Fins were imaged at 1 dpa before injection and at 3 dpa with a Leica MZ16F stereomicroscope. Regenerative fin areas on the injected and uninjected halves were measured using ImageJ and the percentage of regenerative growth was calculated. Statistical significance was determined using one-way ANOVA with multiple comparisons.

### Inhibitor treatment

CB6644 (Cayman Chemical, cat. no. 36125) was dissolved in DMSO to prepare a 10 mM stock concentration and diluted in fish water to a final concentration of 10, 25, and 50 μM, each containing in 1% DMSO. Larvae at 3 dpf with amputated fins were treated with the inhibitor and the treatment was refreshed every 24 hours for 4 consecutive days. At 7 dpf, both control and drug-treated larvae were anaesthetized, and the area of larval fin regenerative growth was measured.

**Supplemental Table S1:** Single-cell analysis of major cell types during zebrafish caudal fin regeneration.

**Supplemental Table S2:** Differentially expressed genes (DEGs) in EpiTag GFP+ cells. Sheet 1: GFP+ vs. GFP-DEGs (log_2(_(Foldchange) >0.6 and adjusted p-value <0.05).

Sheet 2: GFP+ vs. 0 hpa control DEGs (log_2(_(Foldchange) >0.6 and adjusted p-value <0.05).

**Supplemental Table S3:** Clustering of overlapping differentially expressed genes (DEGs) in EpiTag GFP+ cells based on time-course RNA-seq expression profiles.

**Supplemental Table S4:** ATAC-seq differentially accessible regions. Sheet 1: GFP+ vs. GFP- (FDR <0.1 and log2(Fold change) >0.6).

Sheet 2: GFP+ vs. 0 hpa control (FDR <0.01 and log2(Fold change) >0.6).

**Supplemental Table S5:** ATAC-seq footprinting and transcription factor motif enrichment analysis.

Sheet 1: GFP+ vs. GFP- comparison.

Sheet 2: GFP+ vs. 0 hpa control comparison.

**Supplemental Table S6:** Bisulfite-seq differentially methylated regions (DMRs) and nearest gene analysis.

Sheet 1: GFP+ vs. GFP- comparison (mean difference >0.1 and n≥5).

Sheet 2: GFP+ vs. 0 hpa control comparison (mean difference >0.1 and n≥8).

**Supplemental Table S7:** Integrative multi-omics annotation of EpiTag GFP+ differentially expressed genes (DEGs) with associated chromatin accessibility, DNA methylation, cell-type enrichment.

**Supplemental Table S8:** Primer sequences used for RNA probe synthesis.

**Supplemental Table S9:** Percent regenerative growth of zebrafish caudal fins injected with control or gene-targeting Vivo-Morpholinos.

**Supplemental Table S10:** Regenerative fin area (mm^2^) in control-or drug-treated zebrafish larvae.

---

**Figure S1.**
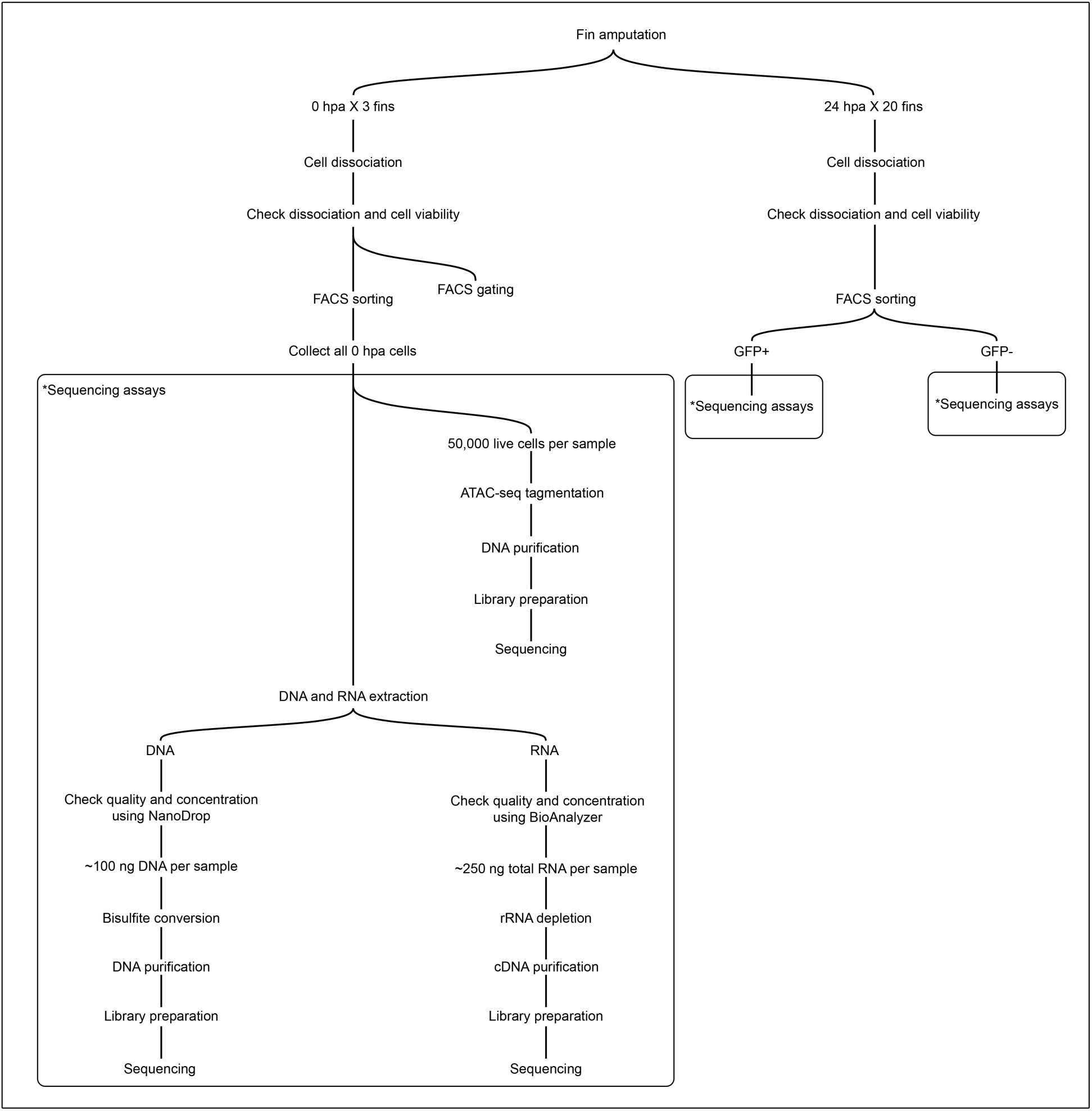
Experimental workflow for EpiTag-based cell sorting and multi-omics profiling. Adult EpiTag zebrafish were subjected to caudal fin amputation, and fin tissues were collected at key regeneration time points. At 0 and 24 hours post amputation (hpa), fins were dissociated into single-cell suspensions. At 24 hpa, cells were FACS-sorted into GFP+ and GFP- populations, while all cells were collected at 0 hpa as a control. These three cell populations were processed in parallel for bulk RNA-seq, ATAC-seq, and bisulfite sequencing.

**Figure S2.**
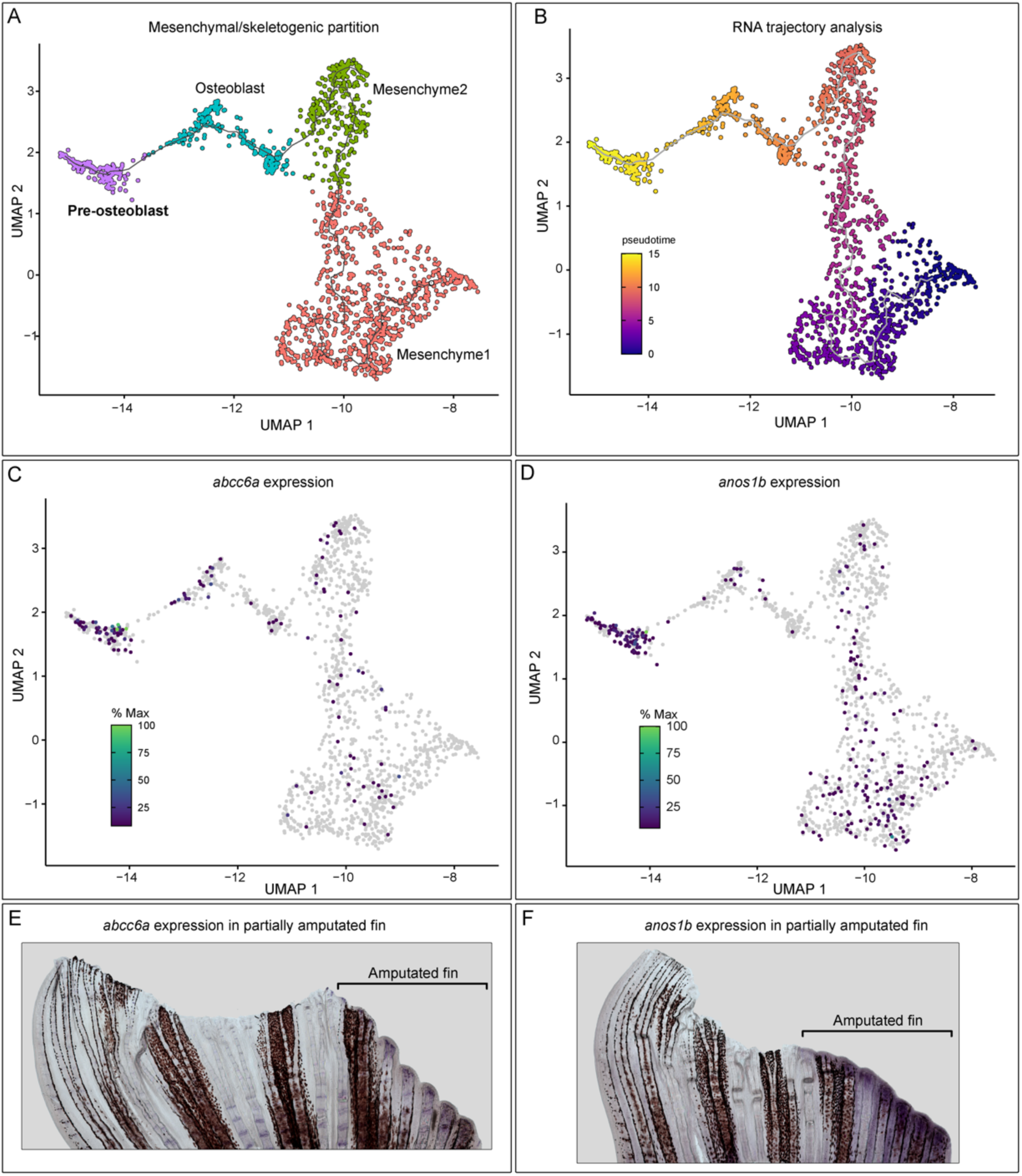
Identification and validation of pre-osteoblast markers. (A) Monocle3 partition of mesenchymal and skeletogenic cell populations, consisting of mesenchyme1, mesenchyme2, osteoblasts, and pre-osteoblasts. (B) RNA trajectory analysis showing that pre-osteoblasts localize downstream of osteoblasts at the terminal end of the differentiation trajectory, supporting a dedifferentiation model. Featureplots of *abcc6a* (C) and *anos1b* (D) reveal strong enrichment in the pre-osteoblast population. WMISH of *abcc6a* (E) and *anos1b* (F) in partially amputated fins.

**Figure S3.**
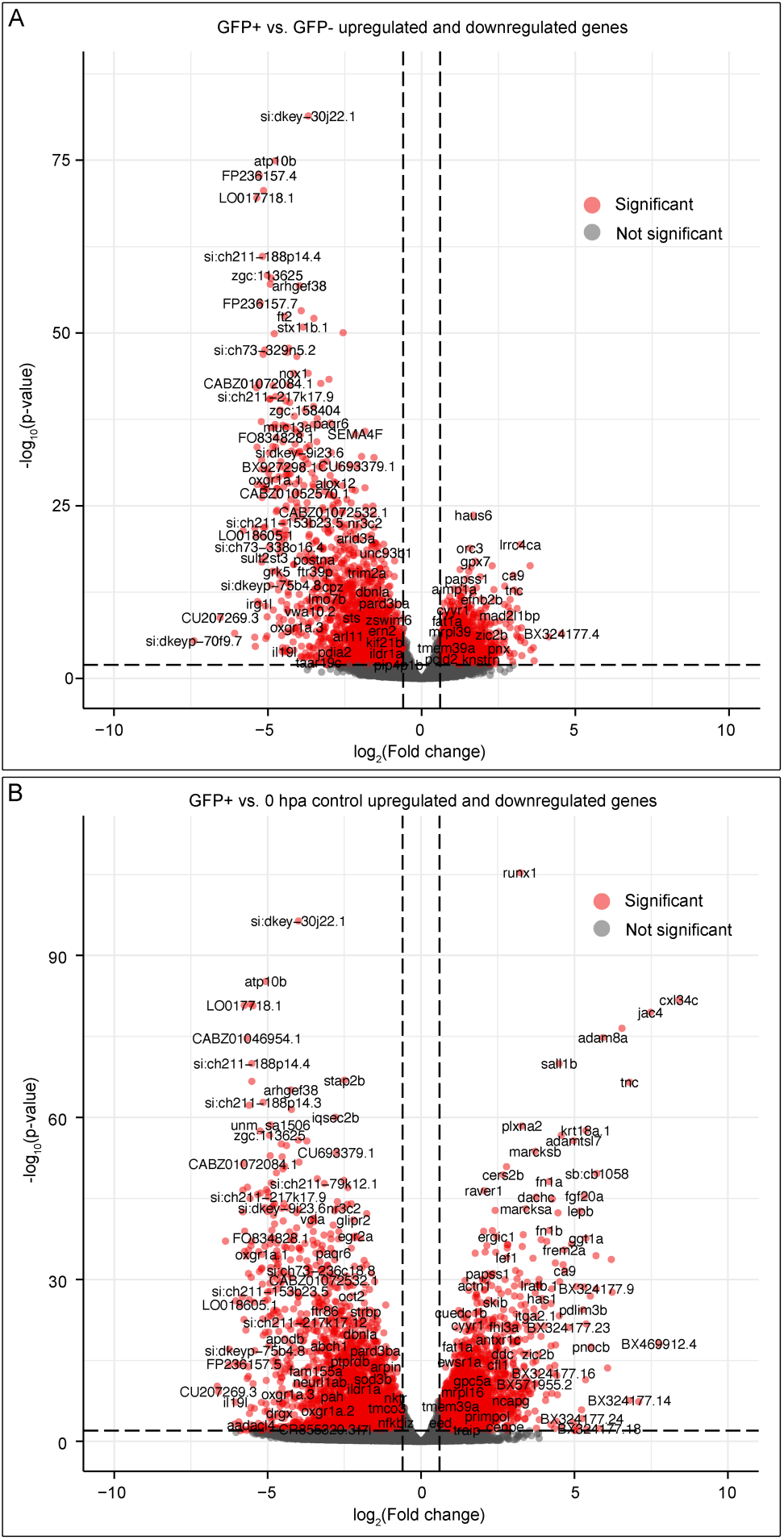
Highly upregulated and downregulated genes in GFP+ cells. (A) Volcano plot showing differentially expressed genes (DEGs) in GFP+ cells compared to GFP- cells isolated at 24 hpa. (B) Volcano plot of DEGs in GFP+ cells compared to 0 hpa controls. For both analysis, genes with adjusted p-value <0.01 and log2(Fold change) <0.6 are highlighted.

**Figure S4.**
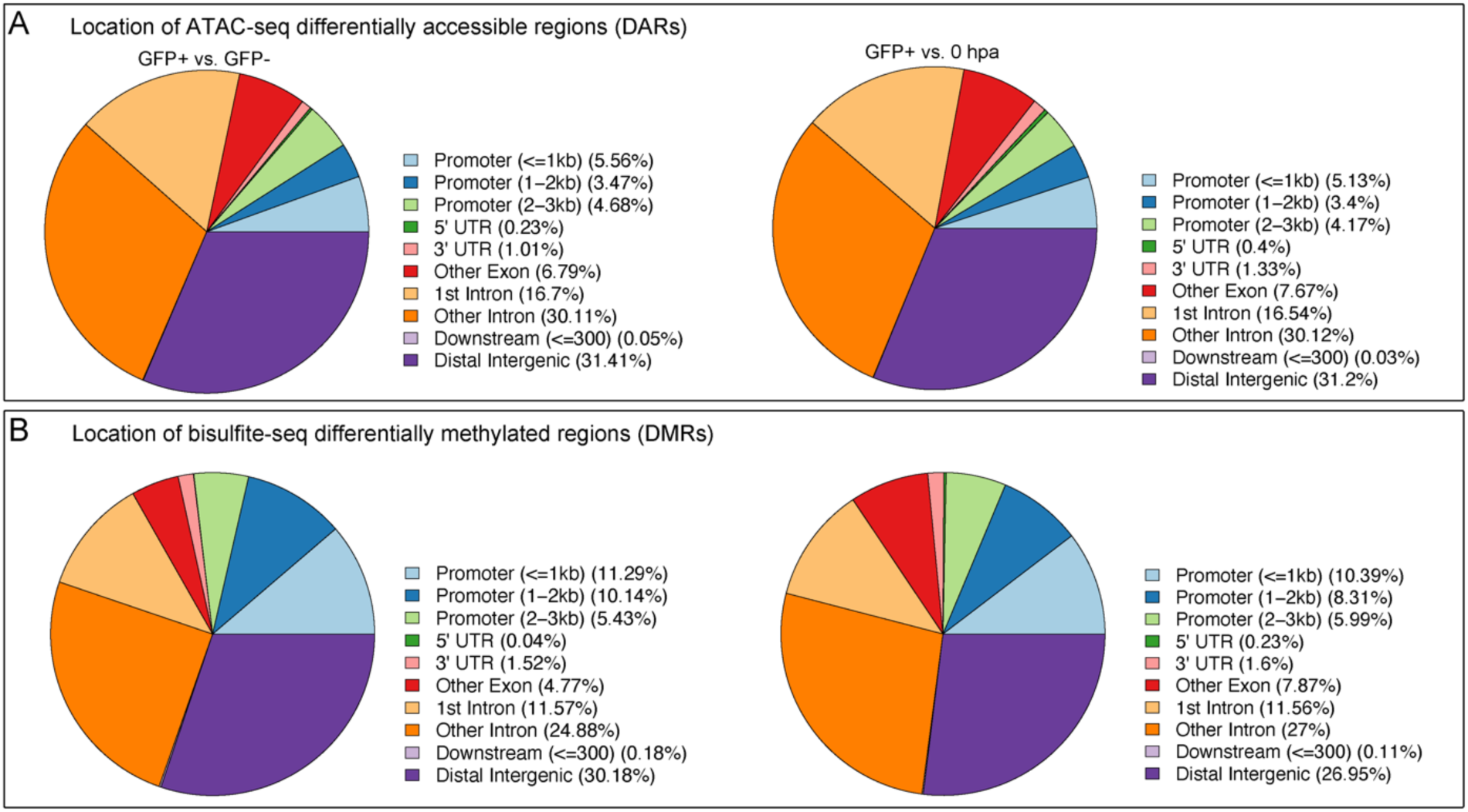
Genomic annotation of differentially accessible regions (DARs) and differentially methylated regions (DMRs) during regeneration. (A) Distribution of ATAC-seq DARs from 24 hpa GFP+ vs. GFP- (left) and 24 hpa GFP+ vs. 0 hpa control (right) comparisons across genomic features. (B) Distribution of bisulfite-seq DMRs from 24 hpa GFP+ vs. GFP- (left) and 24 hpa GFP+ vs. 0 hpa control (right) comparisons across genomic features. For all comparisons, non-coding regions represent the majority of peak-associated regions.

**Figure S5.**
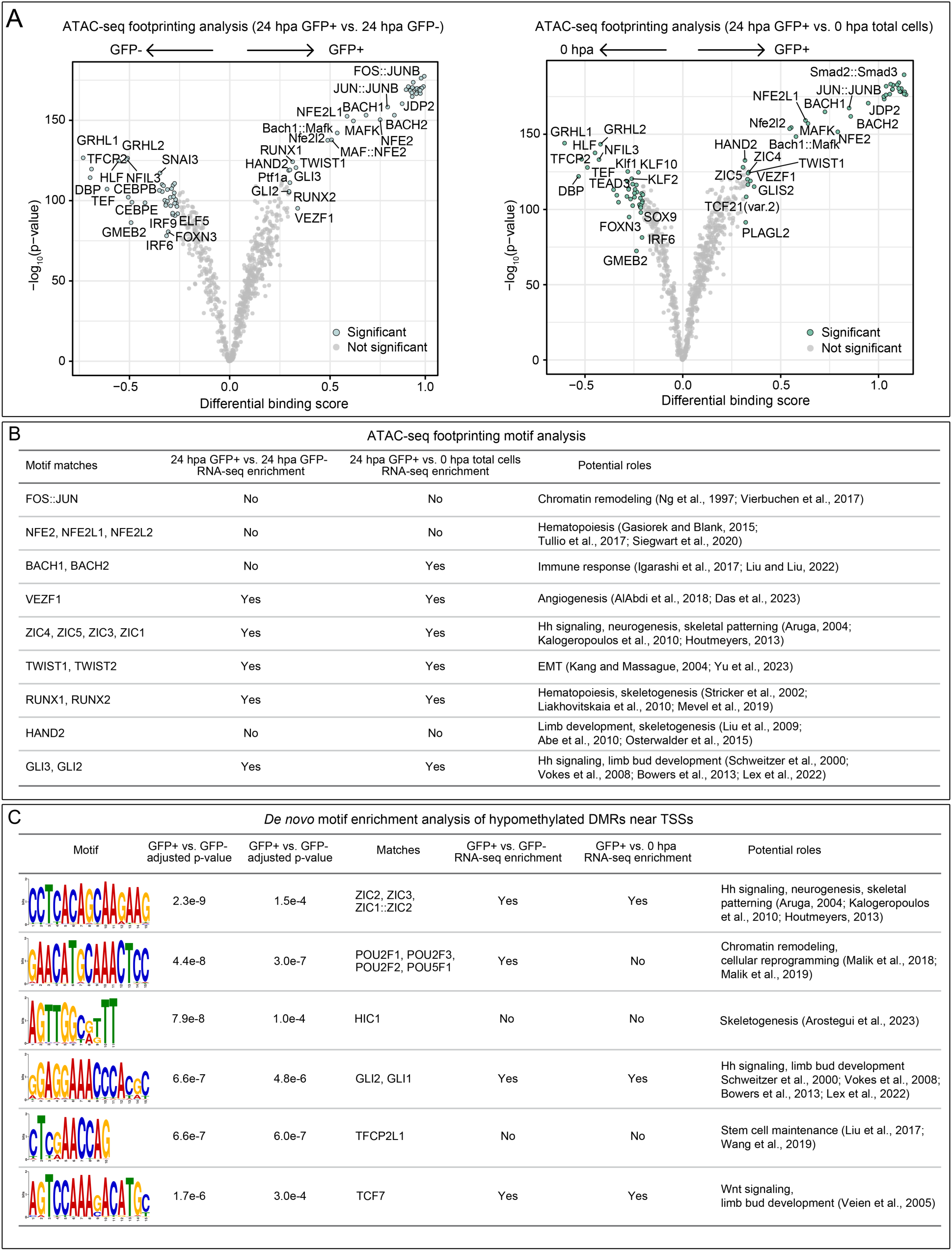
Transcription factors identified by ATAC-seq footprinting and DNA differentially methylated region motif analysis. (A) Volcano plot from TOBIAS footprinting analysis showing transcription factors with differential binding activity between conditions. (B) Summary table of top transcription factors identified by TOBIAS motif enrichment analysis, showing known biological functions with supporting references (Abe et al., 2010; AlAbdi et al., 2018; Arostegui et al., 2023; Aruga, 2004; Bowers et al., 2012; Das et al., 2023; Di Tullio et al., 2017; Gasiorek & Blank, 2015; Houtmeyers et al., 2013; Igarashi et al., 2017; Kalogeropoulos et al., 2010; Kang & Massagué, 2004; Lex et al., 2022; Liakhovitskaia et al., 2010; G. Liu & Liu, 2022; K. Liu et al., 2017; N. Liu et al., 2009; Malik et al., 2018, 2019; Mevel et al., 2019; Ng et al., 1997; Osterwalder et al., 2014; Schweitzer et al., 2000; Siegwart et al., 2020; Stricker et al., 2002; Veien et al., 2005; Vierbuchen et al., 2017; Vokes et al., 2008; Wang et al., 2019; Yu et al., 2023). (C) Summary table of candidate transcription factor motifs identified by STREME de novo motif enrichment analysis of hypomethylated DMRs located near transcription start sites (TSSs), showing known biological functions with supporting references.

**Figure S6.**
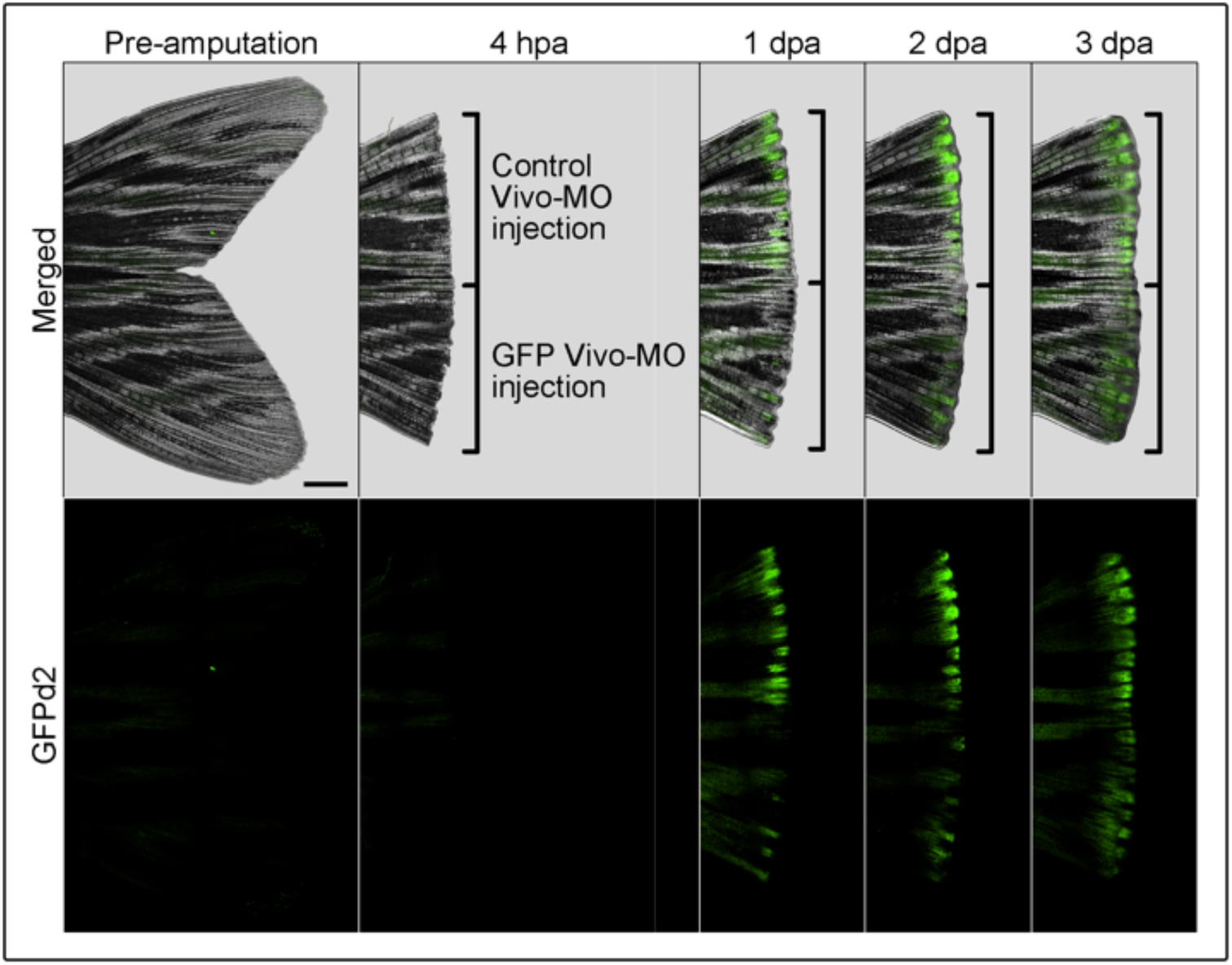
Optimization of GFP Vivo-Morpholino knockdown in regenerating fin. Adult EpiTag fin was injected with GFP Vivo-Morpholino at 4 hpa and imaged at pre-amputation, 4 hpa, 1, 2, and 3 dpa. Efficient knockdown of GFPd2 fluorescence was observed and sustained through 3dpa.

## Notes

### Competing Interest Statement

The authors have declared no competing interest.

### Funding

- Eunice Kennedy Shriver National Institute of Child Health and Human Development: ZIA-HD008977

